# Morpho-physiobiochemical dissection reveals insight into salt-induced differential responses in genetically modified Solanum melongena L. (Bt Brinjal) varieties using an indigenous hydroponic system

**DOI:** 10.1101/2023.05.22.541706

**Authors:** Md. Nahid Hasan, Md. Nazmul Hasan, Md. Nurealam Siddiqui, Md. Arifuzzaman, Mohammad Anwar Hossain, Shamsul H. Prodhan, Md. Ashrafuzzaman

## Abstract

Salinity is a major abiotic constraint of crop production in many countries, including Bangladesh, where a significant amount of cultivable areas are diversely affected by rising salt concentrations. Therefore, it is of paramount importance to figure out the possible solutions to cope with this abiotic stress. So, the development of tolerant genotypes of various crop species can be the best alternative to enhance crop production as well as to improve the livelihoods of poor marginal farmers. With this in mind, the impact of different NaCl levels (50 mM, 100 mM, and 150 mM) on four different Bt Brinjal (Bacillus thuringiensis) genotypes (BARI Bt Begun-1, BARI Bt Begun-2, BARI Bt Begun-3, and BARI Bt Begun-4) was evaluated using morpho-physicochemical analyses at growth, harvesting, and postharvest stages by establishing a new indigenous cost-effective hydroponic system. Our results show that excess salt (> 100 mM) has a detrimental effect on plant growth and development and most of the traits measured across different growth stages. Based on the different measured traits, BARI Bt Begun-1 and BARI Bt Begun-2 varieties outperformed in terms of better morpho-physiological, biochemical, photosynthetic, and antioxidant capacity under salt stress when compared to BARI Bt Begun-3 and BARI Bt Begun-4. Therefore, we conclude that BARI Bt Begun-1 and BARI Bt Begun-2 are moderately tolerant varieties, while BARI Bt Begun-3 and Begun-4 were susceptible varieties to salinity stress. The identified salt-responsive contrasting varieties can serve as valuable genetic materials in comparative genomics for breeding salt-tolerant Brinjal varieties and the newly established hydroponic system could be utilized in translational research programs.

## 1. Introduction

Brinjal (Solanum melongena L.,) commonly known as Eggplant, Aubergine, or Begun (local name in Bangladesh) is a member of the Solanaceae family and is widely cultivated in areas of Asia, Europe, and Africa [23]. It is a perennial herbaceous plant, cultivated year-round and also commercially [11]. There are several stages in the Brinjal life cycle including seed germination (6-8 days), seedling (30-40 days), vegetative (30-60 days), flowering (7-10 days), fruits and maturity (depending on the species) (20-35 days) [11]. For optimal germination of these plants, a moisture content of approximately 60% and a temperature of 30-35°C are necessary for the ideal situation. This globally popular vegetable crop is a leading vegetable in Bangladesh also, and it is the country’s second-most significant vegetable crop in terms of both area and total production [32]. Brinjal production accounted for 4.7 and 9.6 percent of total winter and summer vegetable production in 2018, respectively [33]. After receiving approval from the National Committee on Biosafety (NCB) of Bangladesh in October 2013, four genetically modified Brinjal varieties, namely, Bangladesh Agricultural Research Institute (BARI) Bt Begun-1 (Uttara), BARI Bt Begun-2 (Nayantara), BARI Bt Begun-3 (Kazla), and BARI Bt Begun-4 (Iswardi/ISD006), are now being extensively grown alongside conventional Brinjal varieties [22; 31].

Salt stress is one of the most prominent threats to crop production globally as well as in Bangladesh. It has a significant impact on crop growth and a negative impact on agricultural productivity throughout the world [5; 13; 24]. When plants are exposed to salt stress salts, it affects their internal physiological processes, molecular mechanisms, and macromolecule chemical functions, resulting in lower yields [4; 27]. Soil salinity is known to inhibit plant growth by causing osmotic stress, which is followed by ion toxicity [20; 29]. Since 1973, Bangladesh’s overall salinity-affected land area has grown by 26.7% from 83.3 million ha to 105.6 million ha in 2009. Salt-affected soils limit the amount of water and nutrients that plants can absorb from the soil, which eventually causes an imbalance in osmotic potential, and ionic equilibrium and has an impact on plant’s physiology, growth, and development [24] and reduces yields significantly [5; 14]. This is because of the toxic ion concentrations in the soil that have been shown to impede plant development by inducing osmotic stress [20; 29]. Osmotic stress causes a variety of physiological changes in the early stages of salinity stress, including membrane disruption, impaired ability to detoxify reactive oxygen species (ROS), nutrient imbalance, decreased photosynthetic activity, differences in antioxidant enzymes, and a decrease in stomatal aperture [24; 29]. Moreover, in the presence of salt stress, reactive oxygen species (ROS) such as superoxide, hydroxyl radicals, and hydrogen peroxide (H_2_O_2_) are formed. ROS generation generated by salinity can cause oxidative damage to a variety of cellular components, including proteins, lipids, and DNA, resulting in the disruption of key physiological activities in plants [16].

Therefore, salt stress should be addressed appropriately in order to ensure agricultural sustainability and the continuation of food production. Conventional plant breeding procedures are used to improve the tolerance of plants against salinity, which are time-consuming and labor-intensive, and they rely on genetic variability that already exists in the populations [18]. At the moment, the majority of efforts are focused on the genetic transformation of plants, with traditional physiological treatments being abandoned. The employment of physiological and biochemical selection criteria can increase the resolution of the mechanistic basis of salt stress responses. Fewer new varieties with higher salt tolerance have been developed by selection for agronomic traits until now, despite the fact that our knowledge of physiological responses has expanded dramatically during the same period of time. One technique for dealing with the problem of salinity is to choose genotypes that have the ability to withstand an excessive amount of salt. For marginal farmers in the coastal region, a high-producing salt-tolerant crop variety is the ultimate dream. For this, the current study was conducted with the aim to test the suitability and feasibility of Bt Brinjal genotypes by assessing an integrated approach combining morphology, physiology, and biochemical attributes. Besides this, we aimed to establish a cost-effective indigenous hydroponics system for plant cultivation and to explore the effects of salt on different growth, physiological and biochemical parameters of four Bt Brinjal varieties (BARI Bt Begun-1, BARI Bt Begun-2, BARI Bt Begun-3, and BARI Bt Begun-4).

**Figure 1.**
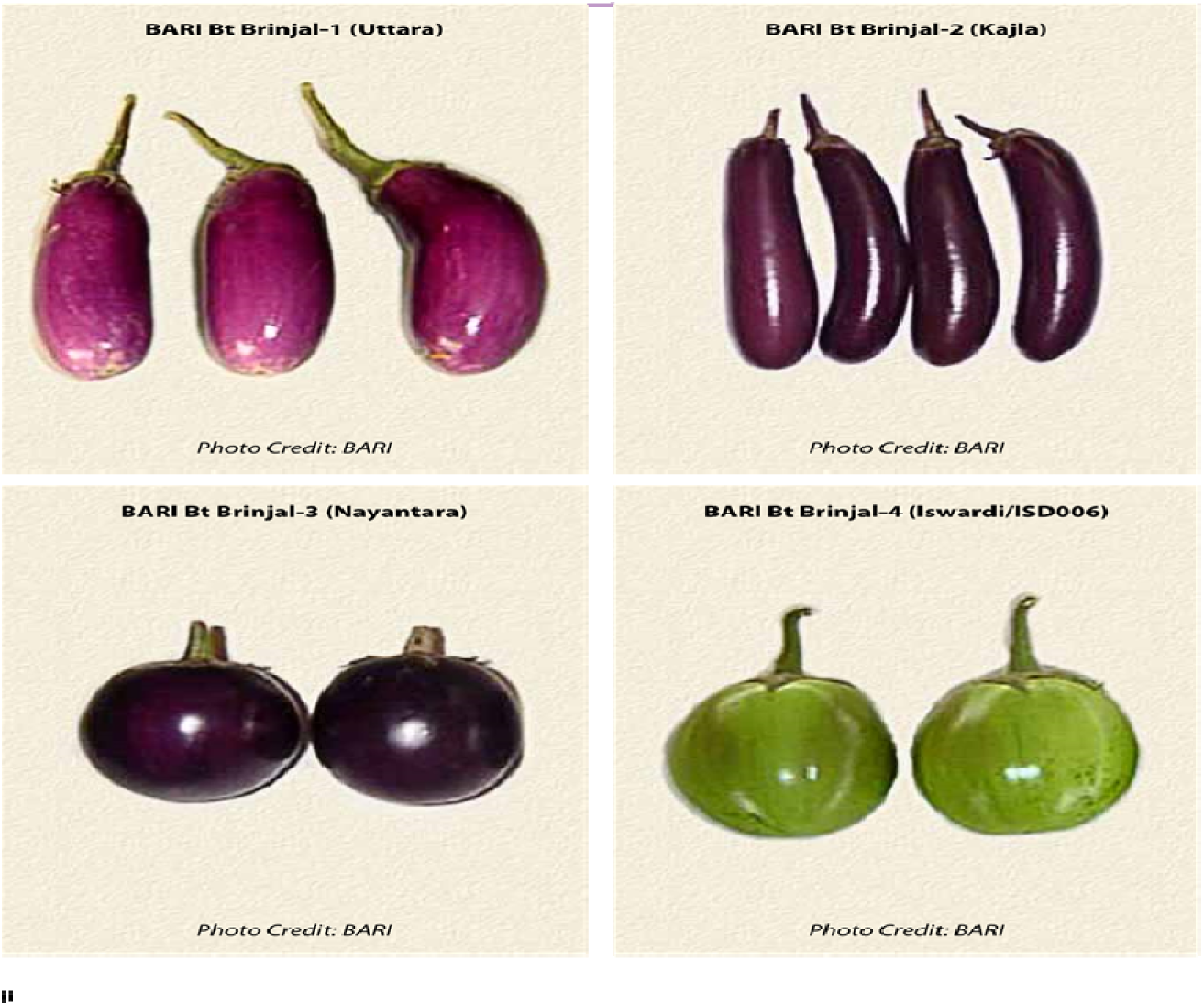
Four cultivatedBt Brinjal varieties [12] 2. Materials and methods

## 2. Materials and methods

### 2.1 Plant materials and growth conditions

Four Bt Brinjal varieties, namely BARI Bt Begun-1(Uttara), BARI BtBegun-2 (Kajla), BARI Bt Begun-3 (Nayantara), and BARI Bt Begun-4 (ISD006), developed by Bangladesh Agricultural Research Institute (BARI), were screened for their salt tolerance levels at the seedling stage. A controlled environment at the Greenhouse of Plant Genetic Engineering Laboratory, Department of Genetic Engineering and Biotechnology, Shahjalal University of Science and Technology, Sylhet-3114, was maintained for this experiment. The average measured temperature was 26 ±2°C, 16 hours of light and 8 hours of dark condition were maintained, and the average light intensity ranged between 1500 and 2000 lux. The experiment was conducted in large rectangular-shaped containers (24-inch length×16-inch width × 13-inch height), which can hold 68 L of water [6]. A plastic lid with 40 holes attached to plastic pipes was used to grow the plants. The radius of the hole was 1.5 centimeters. Styrofoam sheets were used to hold the plants (Figure 2). With three replications, the experiment was conducted in a randomized complete block design.

**Figure 2.**
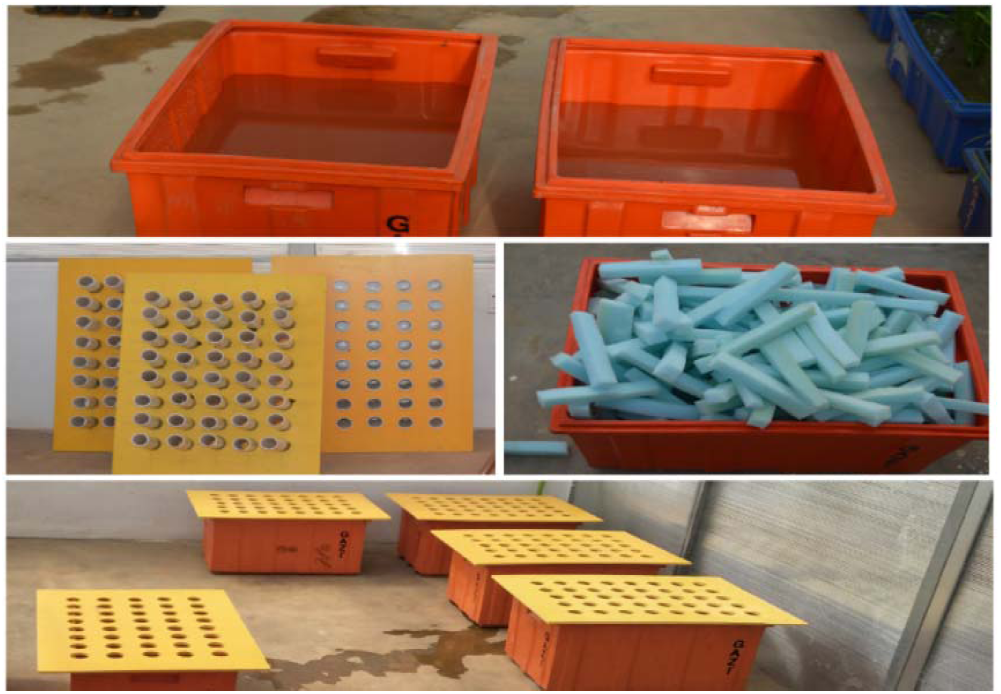
Experimental setup of the hydroponics system

### 2.2 Seed germination and seedling establishment in the hydroponics system

The seed germination was done in a germination tray. We used cocopeat for the germination of seeds. At first, the seeds were dried in sunlight for around 6 hours. After drying, the seeds were soaked in water for about 4 hours. Then seeds were sowed into the cocopeat on the germination tray. The germination process took around seven days to become two leaf stages. The process took a total of around 32 days to become the plant to the seedling stage. During those time periods, the moisture content and light intensity were carefully maintained. The seedlings were then transferred into 68-L plastic containers filled with half-strength modified Yoshida nutrition solution [36] (Supplementary Figure 1). Seedlings were kept in the half-strength solution for one week, and after that, the solution was replaced by a full-strength solution subsequently every week for the rest of the experiment. The pH was adjusted twice a week to 5.5 and maintained throughout the experiment.

Liquid nutrient media was used to grow the Brinjal seedlings. In accordance with Yoshida (1976), a modified nutrient solution was made by following changed compositions, and it is comprised of six stock solutions (five for major elements and one for all micro-elements) (Supplementary Table 1). At first, 5 Lof six stock solutions were prepared and stored in the dark brown colored glass bottle and kept at room temperature. After that, the presence of mineral precipitation or a change in the covalence of elements such as iron or copper in the solution was investigated. The storage containers were maintained air and light-tight for long storage. The working solution was prepared as described by [36] and [6]. Since our container carried 68 Lof distilled water, we added each stock solution in the amount of 85 mL in one container to prepare full strength working solution. With constant stirring, 1N NaOH and 1N HCl were used in the working solution to maintain the pH level to 5.5. A decrease in solution volume and a shift in pH are both caused by evaporation and transpiration. After every three days, the volume was brought back to the desired level, and the pH was adjusted twice a week to 5.5.

### 2.3 Treatment of salt stress

The treatments were comprised of four salinity levels, 0 mM (control), 50 mM, 100 mM, and 150 mM NaCl. After 7 days of growth of plants in the full-strength nutrient solution, the NaCl treatment was administered in the setup. We applied the salt treatment at the 4-leaf stage. As a precaution, a gradual 50 mM NaCl solution was added to the salt containers each day to attain a final salt concentration of 150 mM NaCl to avoid the sudden osmotic shock in the plants. The electrical conductivity of the salt-treated nutrient solution was checked and maintained at the desired level of salt concentration with electrical conductivity (EC) meter (Hanna Instruments, HI98303), and the treatment continued for 21 days up to enough symptoms appeared.

### 2.4 Determination of phenotypic and physiological traits

The Brinjal plants were treated in their respective salt concentration for 21 days. Salt Evaluation Score (SES) and the number of leaves/plants were determined visually only before harvesting. A modified salt scoring system (0 to 10) was used to quantify visible leaf symptoms induced by NaCl stress in an average of the fully expanded lower to upper leaves of each plant [6; 15]. In the scale, the criteria were as follows: 1 indicates no stress damage symptoms in any part of the leaf, whereas 3, 5, 7, and 9 define leaves with approximately 30%, 50%, 70%, and 90% damage due to the salt stress, respectively. The root length (cm), shoot length (cm), root fresh weight (gm), shoot fresh weight (gm), root dry weight (gm), and shoot dry weight (gm) were measured after harvesting. For weighing an electronic balance was used to record fresh weight immediately upon harvest to minimize evaporation. Shoot and root samples were oven-dried at 60 °C for at least 72 hr and weighed. Leaf samples for biochemical analysis were collected using liquid nitrogen and stored at freeze for the analysis.

### 2.5 Biochemical analyses

#### 2.5.1 Measurement of total chlorophyll content and carotenoid content

0.1 g of leaf sample was taken and used to measure the total chlorophyll and carotenoid contents for each sample with 3 replications, according to [3]. Three measurements were recorded for each sample and then averaged. The amounts of chlorophylls and carotenoids were estimated using the following equations [21]:

Total chlorophyll (mg/g fresh weight) = [20.2 (A645) + 8.02 (A663)] × [V/ (1000 W)]

Carotenoid (mg/g fresh weight) = [A480 + (0.114(A663)) – (0.638(A645))] × [V/ (1000 W)]

Where, A = Absorbance at specific wavelengths

V = Final volume of chlorophyll extract in 80% acetone

W = Fresh weight of tissue extracted.

#### 2.5.2 Measurement of proline content

Proline content was estimated by the procedure described by Bates et al.(1973) [9] 0.1 g fresh weight of leaf tissues was used for each sample with three replications. We used a cuvette and spectrophotometer to measure color intensity at 520 nm absorbance. We also used pure standard proline in a similar way, and a standard curve was prepared. Then test sample was calculated from the standard curve[10].

The proline content on a fresh-weight-basis was expressed as follows:

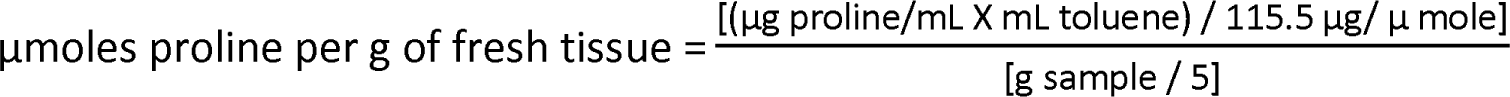

Here, 115.5 is the molecular weight of the proline.

#### 2.5.3 Determination of antioxidant enzyme activities

At first, enzyme extracts were produced to measure the activity of catalase (CAT) and peroxidase (POD). For enzyme extract preparation, 0.1 g fresh weight of leaf tissues was homogenized in 5 ml of 10 mM potassium phosphate buffer (pH 7.0) containing 4% (w/v) PVP. In order to assess enzyme activity, the supernatants from the homogenate were immediately centrifuged at 12,000 rpm for 15 minutes at 4°C. The resulting enzyme extract was then used for the assay of enzyme activities, viz. catalase (CAT) and peroxidase (POD). The activity of the CAT and the activity of the POD was measured using the procedure published by Velikova et al. (2000) [35], with a few minor adjustments. We used a spectrophotometer to measure the absorbance. The activity of the CAT enzyme was calculated using the extinction coefficient, e of 40 mM^-1^cm^-1^ 0.04 M^-1^cm^-1,^ and the extinction coefficient, e of 26.6 mM^-1^cm^-1^for POD activity calculation. We used the following equation for calculating the CATand POD activity:

Enzyme activity (Units/L) = (ΔAbs × Total assay volume) / (Δt x ε x l x Enzyme sample volume) Where Δt is the time of incubation (min), ΔAbs is the change in absorbance, ε is the extinction coefficient of substrates in units of M^-1^cm^-1^), and l is the cuvette diameter (1cm). Enzyme activity (Unit) was defined as the amount of enzyme that oxidized 1μmol of substrate/min [1].

#### 2.5.4 Measurement of hydrogen peroxide content

Determination of H_2_O_2_ content in plant tissues was done following [35]. In our experiment, we used 0.1 g fresh weight of leaf tissues for each sample with three replications. The content of H_2_O_2_ was calculated using Beer’s law A=€bc, where A is absorbance, e is molar extinction coefficient (€€ = 0.28 μM cm^-1^), b is the path length of the sample, that is, the path length of the cuvette in which the sample is contained, and c is the concentration of the compound in solution, expressed in mol L^-1^ [25]. In our equation, the unit of c is μmol/gFW.

#### 2.5.5 Quantitative analysis of total anthocyanin content

Total anthocyanin content was analyzed by the procedure of [2]. The calculation was done using the equation given below:

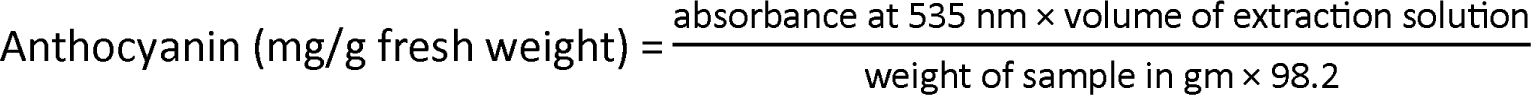

The concentration was calculated according to Beer’s law A=€bc where A is absorbance, e is molar extinction coefficient, b is the path length of the sample, that is, the path length of the cuvette in which the sample is contained, and c is the concentration of the compound in solution, expressed in mol L^-1^ [25].

### 2.6 Statistical analysis

The experiment was conducted in a randomized complete block design with three replications. All statistical analyses including two-way analysis of variance (ANOVA) were performed using SigmaPlot version 12.5 (Systat Software, San Jose, CA). All pairwise mean comparison was performed using Tukeytest and P-values less than 0.05 were considered as significant.

## 3. Results

### 3.1 Phenotypic changes under different salinity levels

The four Bt Brinjal varieties grown as a control (without any salt treatments) had normal phenotypic growth and were healthy in all aspects. But plant development was affected when the plants were exposed to varying salt concentrations. The severity of the damage was proportional to the level of salinity strength given. The plants grown in 50 mM NaCl concentration of salt stress exhibited similar or sometimes a slight reduction in growth parameters compared to control plants. In contrast, the plants treated with 100 mM and 150 mM NaCl stress showed a greater and clear drop in growth performance and appearance after 21 days of treatment in the hydroponic system (Figure 3).

**Figure 3.**
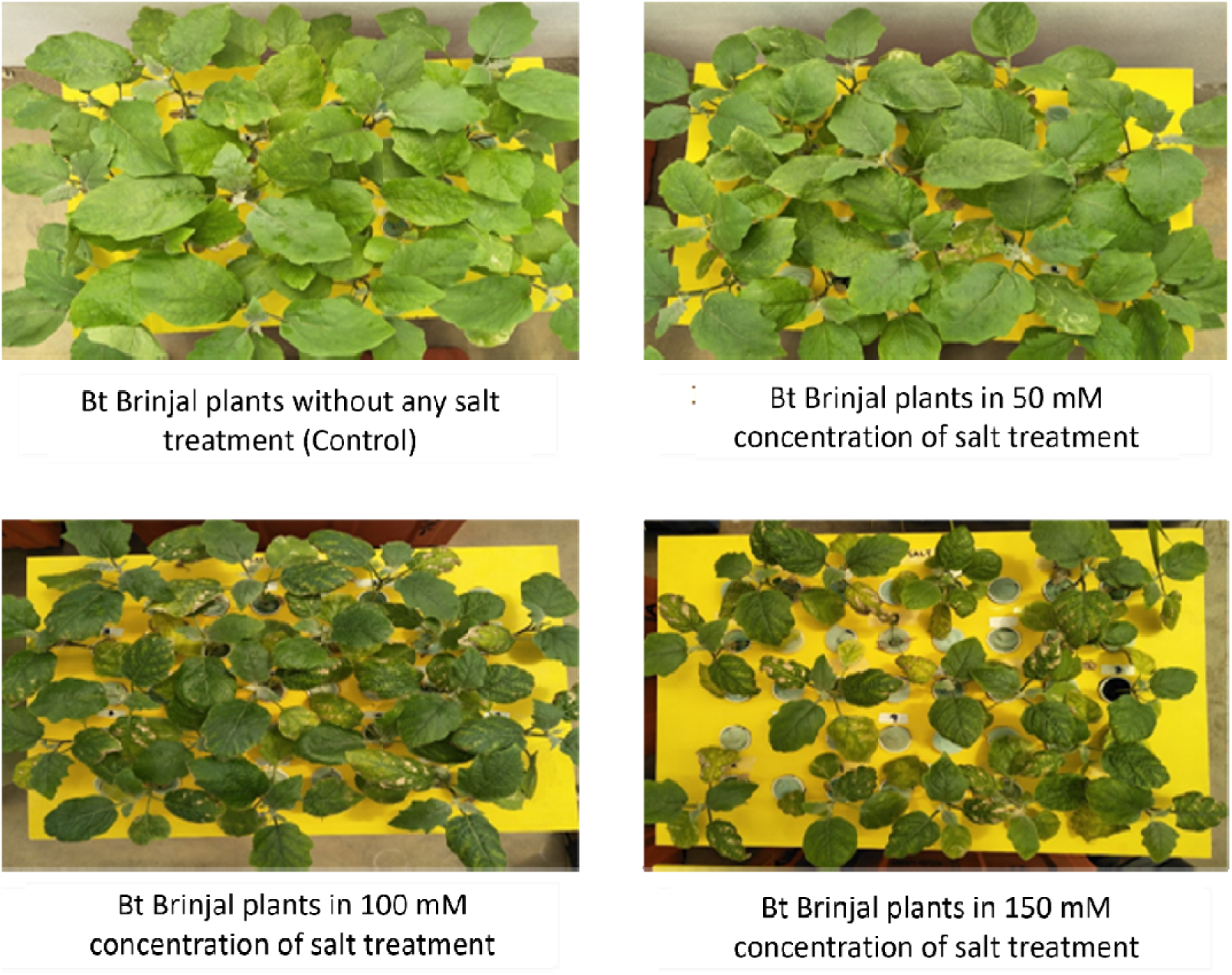
Four Bt Brinjal varieties under the different salt concentrations

Within each genotype, a clear and significant difference was seen in different concentrations of salt treatment, such as 0 mM (control), 50 mM, 100 mM, and 150 mM. A decrease in growth performance and an increase in leaf damage was also seen in 100 mM and 150 mM concentration of salt treatment compared to control and 50 mM concentration of salt treatment (Supplementary Figure 2). All Bt Brinjal varieties’ growth and development performances against different salt treatments were assessed thus revealed Bt-1 (Bt Begun-1) and Bt-2 (BARI Bt Begun-2) varieties showed better morphological and growth patterns under salt treatments when compared to Bt-3 (BARI Bt Begun-3) and Bt-4 (BARI Bt Begun-4) (Supplementary Figure 3).

### 3.2 Morpho-physiological characterization

In a total 8 traits were measured for the Morpho-physiological characterization in which most of them showed significant responses to salt stress in treatment, genotype, and their interaction (Table 1). On the other hand, when averaged over all four genotypes’ mean performance, significant differences were observed in 100 mM and 150 mM NaCl concentrations compared to the control (Table 1). The root length and shoot length in all genotypes of Brinjal seedlings were significantly reduced in 100 mM and 150 mM salt concentrations compared to the control (Figure 4). Similarly, root fresh weight, shoot fresh weight, root dry weight, and shoot dry weight also decreased significantly at 100 mM and 150 mM NaCl compared to the control (Figure 5). All four Bt Brinjal varieties were significantly affected by 100 mM and 150 mM NaCl levels based on salt injury score (Figure 6. A). Besides this, the number of leaves in Bt Brinjal varieties was significantly reduced in 100 mM and 150 mM NaCl concentrations compared to control and 50 mM NaCl level in all genotypes (Figure 6. B).

**Table 1.** Statistical analysis and treatment mean values (two-way ANOVA using SigmaPlot 12.5) of measured growth and physiological parameters from four different Bt Brinjal varieties exposed to control and salinity treatments.

| Traits | ANOVA results (Pr> F) |  |  | LS means (Treatment) |  |  |  |
| --- | --- | --- | --- | --- | --- | --- | --- |
|  | Treatment<br>(T) | Genotype<br>(G) | T×G Interaction | Control | 50 mMNaCl | 100 mMNaCl | 150 mMNaCl |
| Salt evaluation score (SES) | <.0001 | <.0001 | 0.047 | n.d. | 1.75<sup>b</sup> | 5.0<sup>a</sup> | 5.9<sup>a</sup> |
| Number of leaves/plant | <.0001 | <.0001 | 0.011 | 9.7<sup>a</sup> | 7.8<sup>a</sup> | 6.8<sup>b</sup> | 6.4<sup>b</sup> |
| Shoot fresh weight (gm) | <.0001 | <.0001 | <.0001 | 15.2<sup>a</sup> | 12.8<sup>a</sup> | 9.3<sup>b</sup> | 7.5<sup>b</sup> |
| Root fresh weight (gm) | <0.001 | <0.001 | <.0001 | 4.7<sup>a</sup> | 4.2<sup>a</sup> | 3.4<sup>b</sup> | 2.8<sup>b</sup> |
| Shoot length (cm) | <0.001 | <0.001 | 0.015 | 35.4<sup>a</sup> | 31.5<sup>a</sup> | 25.9<sup>b</sup> | 22.0<sup>b</sup> |
| Root length (cm) | <.0001 | <.0001 | <.0001 | 38.0<sup>a</sup> | 35.0<sup>a</sup> | 26.9<sup>b</sup> | 22.7<sup>b</sup> |
| Shoot dry weight (gm) | <0.001 | 0.301 | 0.005 | 1.90<sup>a</sup> | 1.50<sup>a</sup> | 1.21<sup>b</sup> | 0.94<sup>b</sup> |
| Root dry weight (gm) | <0.001 | <0.001 | 0.649 | 0.352<sup>a</sup> | 0.253<sup>a</sup> | 0.167<sup>b</sup> | 0.123<sup>b</sup> |

LS means = least square means. LS mean values not sharing the same superscript letter differ significantly from each other at $P<0.05$ by Tukey-multiple comparison, and n=12. In the table, n.d. means not determined.

**Figure 4.**
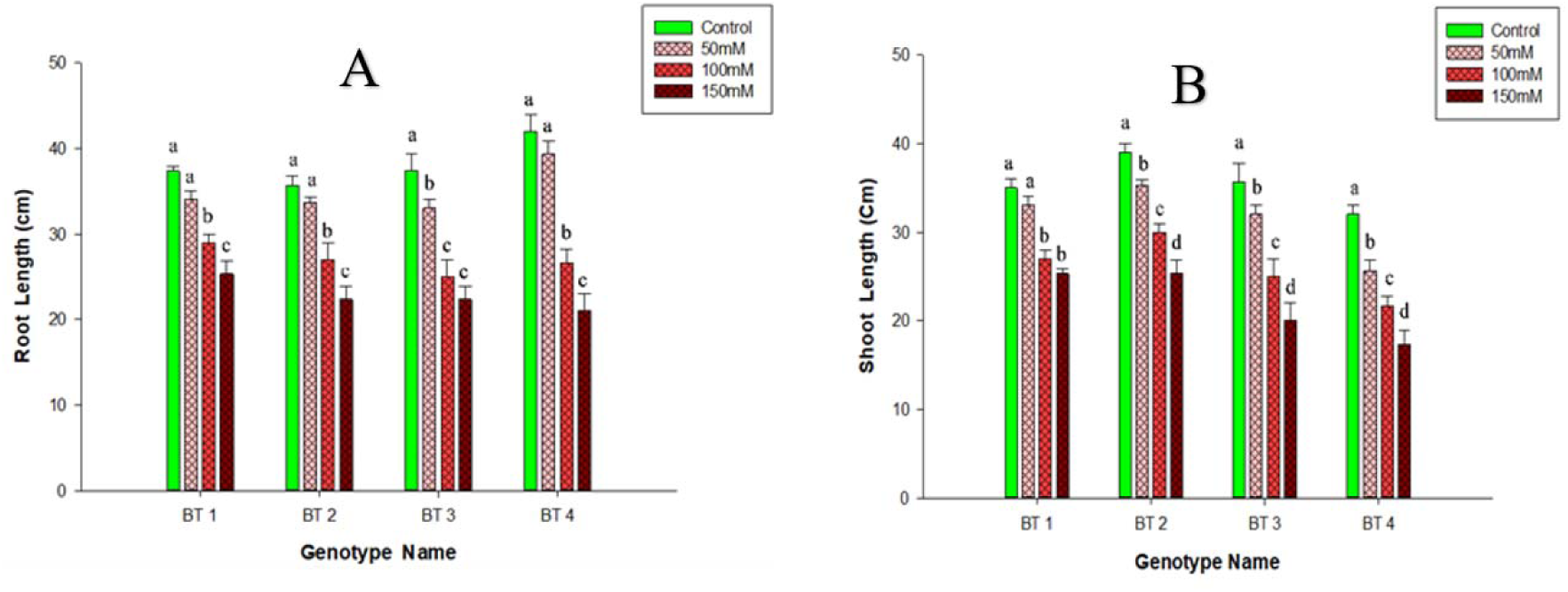
Root length (cm) and shoot length (cm) of four Bt Brinjal varieties in control and three different salinity levels (50 mM, 100 mM, and 150 mMNaClconcentrations). Y-axis represents the Root length (cm) and the shoot length (cm) in Figure 4. A and 4. B, respectively, and each bar indicates mean value ± standard errors (n=3). The upper letter in each bar indicates a pair-wise comparison (P<0.05) within the genotype. The upper letter in the bar not sharing the same letter differs significantly from each other. The genotype name illustrates Bt-1 (Bt Begun-1), Bt-2 (BARI Bt Begun-2), Bt-3 (BARI Bt Begun-3), and Bt-4 (BARI Bt Begun-4).

**Figure 5.**
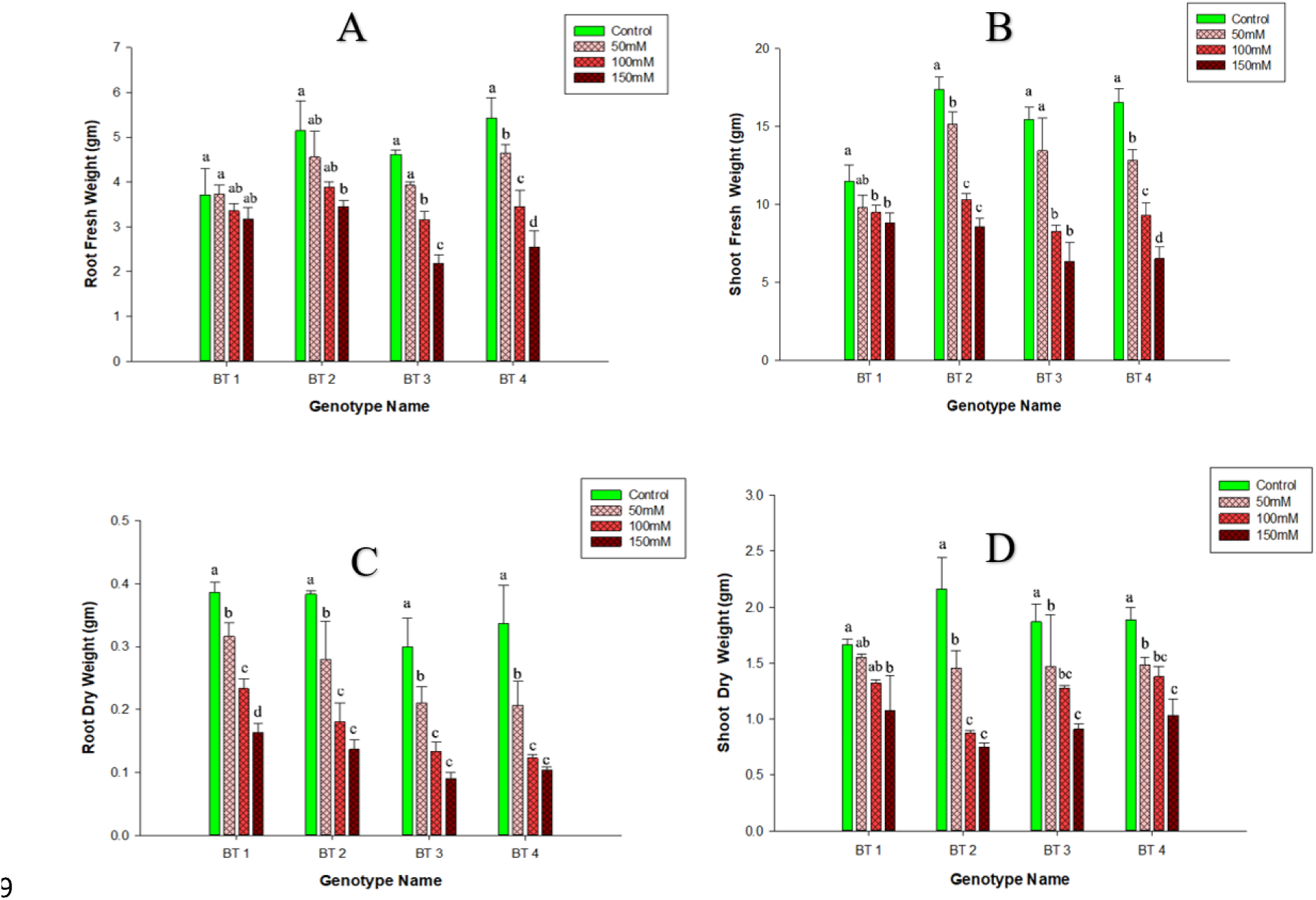
Root fresh weight (gm) (A), shoot fresh weight (gm) (B), root dry weight (gm) (C), Shoot dry weight (gm) (D) of four BT Brinjal varieties in control and three different salinity levels (50 mM, 100 mM, and 150 mM NaCl concentrations). Y-axis represents weights, and each bar indicates mean value ± standard errors (n=3). The upper letter in each bar indicates a pair-wise comparison (P<0.05) within the genotype. The upper letter in the bar not sharing the same letter differs significantly from each other. The genotype name illustrates Bt-1 (Bt Begun-1), Bt-2 (BARI Bt Begun-2), Bt-3 (BARI Bt Begun-3), and Bt-4 (BARI Bt Begun-4).

**Figure 6.**
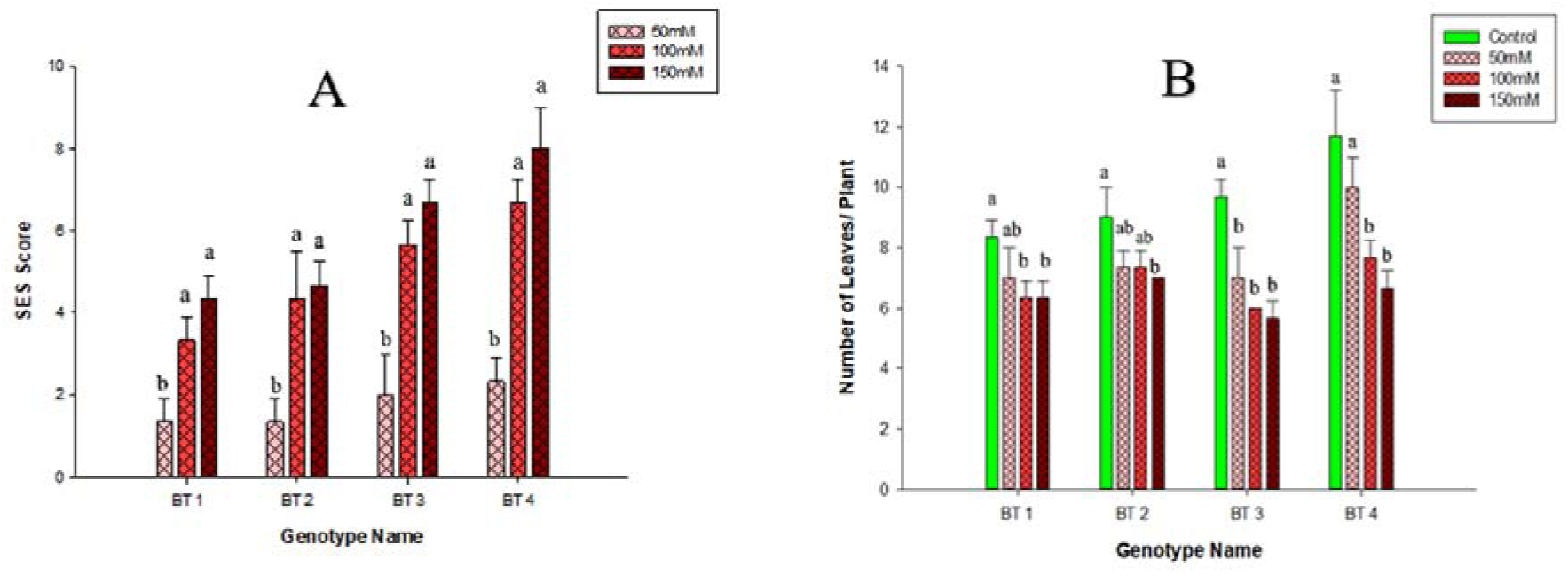
Salt evaluation score (SES) and numbers of leaves/plant of four BT Brinjal varieties in control and three different salinity levels (50 mM, 100 mM, and 150 mMNaClconcentrations). Y-axis represents the SES score (6. A) and the number of leaves/plants (6. B), and each bar indicates the mean value ± standard errors (n=3). The upper letter in each bar indicates a pair-wise comparison (P<0.05) within the genotype. The upper letter in the bar not sharing the same letter differs significantly from each other. The genotype name illustrates Bt-1 (Bt Begun-1), Bt-2 (BARI Bt Begun-2), Bt-3 (BARI Bt Begun-3), and Bt-4 (BARI Bt Begun-4).

### 3.3 Biochemical characterization

For the biochemical characterization, 8 different traits were measured and most of them significantly differ in response to salt stress in treatment, genotype, and their interaction (Table 2). Moreover, significant differences were observed in 100 mM and 150 mM NaCl concentrations compared with control (without NaCl) when averaged over all four genotypes’ mean performance (Table 2).

**Table 2.** Statistical analysis and treatment mean values (two-way ANOVA using SigmaPlot 12.5) of measured biochemical parameters from four different Bt Brinjal varieties exposed to control and salinity treatments.

| Traits | ANOVA results (Pr> F) |  |  | LS means (Treatment) |  |  |  |
| --- | --- | --- | --- | --- | --- | --- | --- |
|  | Treatment | Genotype | Interaction | Control | 50 mMNaCl | 100 mMNaCl | 150 mMNaCl |
| Total chlorophyll content (mg/gm) | <0.001 | 0.051 | 0.085 | 1.297<sup>a</sup> | 1.147<sup>a</sup> | 0.942<sup>b</sup> | 0.833<sup>b</sup> |
| Total carotenoid content (mg/gm) | <0.001 | <0.001 | 0.017 | 0.006<sup>a</sup> | 0.005<sup>a</sup> | 0.003<sup>b</sup> | 0.002<sup>b</sup> |
| Proline content (mmoles/g FW) | <0.001 | <0.001 | 0.093 | 5.997<sup>a</sup> | 8.927<sup>a</sup> | 13.072<sup>b</sup> | 14.896<sup>b</sup> |
| Anthocyanin Content (mg/g) | <0.001 | <0.001 | 0.029 | 0.129<sup>a</sup> | 0.147<sup>a</sup> | 0.170<sup>b</sup> | 0.210<sup>b</sup> |
| Hydrogen peroxide content<br>(μmol/gFW) | <0.001 | <0.001 | 0.816 | 0.373<sup>a</sup> | 0.390<sup>a</sup> | 0.419<sup>b</sup> | 0.451<sup>b</sup> |
| MDA content (mmoles/g FW) | <0.001 | <0.001 | 0.009 | 0.246<sup>a</sup> | 0.306<sup>a</sup> | 0.380<sup>a</sup> | 0.583<sup>b</sup> |
| Catalase activity (Unit/L) | <0.001 | <0.001 | 0.095 | 65.704<sup>a</sup> | 73.058<sup>a</sup> | 77.515<sup>b</sup> | 85.780<sup>b</sup> |
| Peroxidase activity (Unit/L) | <0.001 | <0.001 | 0.068 | 8.748<sup>a</sup> | 14.212<sup>b</sup> | 17.835<sup>b</sup> | 24.051<sup>c</sup> |

LS means = least square means. LS mean values not sharing the same superscript letter differ significantly from each other at $P<0.05$ by Tukey-multiple comparison, n=12

#### 3.3.1 Photosynthetic pigments

Leaf pigments were measured to estimate the leaf health status in the experimental plants. Total leaf chlorophyll and leaf carotenoid contents were decreased significantly in high salt 100 mM and 150 mM concentrations compared to the control (Figure 7. A, B). Interestingly, Bt-1 (BARI Bt Begun-1) and Bt-2 (BARI Bt Begun-2) showed less sensitivity to 50 mM, 100 mM, and 150 mM concentrations of salt treatment because the decrease of chlorophyll content was not significant but the other two genotypes Bt-3 (BARI Bt Begun-3), and Bt-4 (BARI Bt Begun-4) showed a significant decrease in chlorophyll content (Figure 7. A). Along with this, Bt-1 and Bt-2 showed slight tolerance in terms of carotenoid contents compared to the other two Bt genotypes (Figure 7. B).

**Figure 7.**
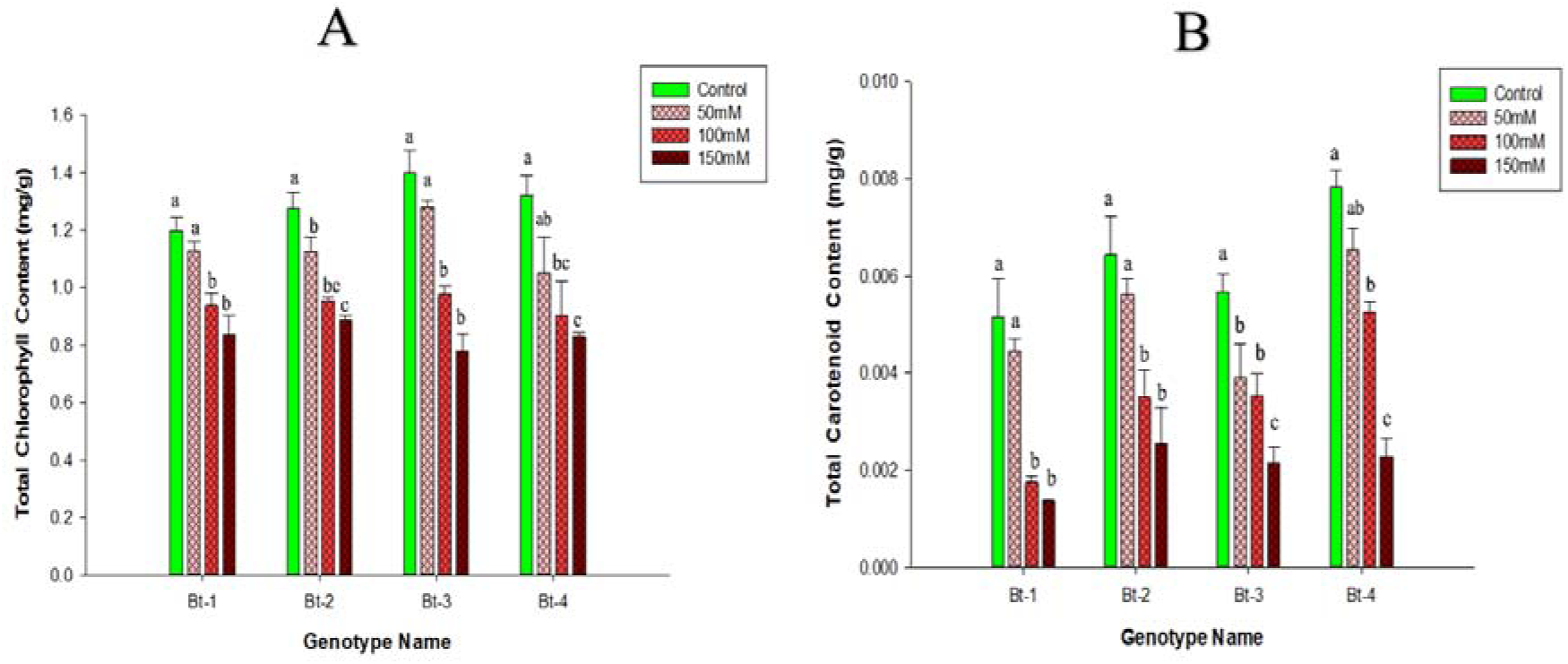
Total chlorophyll content and total carotenoid content of four Bt Brinjal varieties in control and three different salinity levels (50 mM, 100 mM, and 150 mM NaCl concentrations). Y-axis represents the total chlorophyll content (mg/g) (7. A) and total carotenoid content (mg/g) (7. B), respectively. Each bar indicates mean value ± standard errors (n=3). The upper letter in each bar indicates a pair-wise comparison (P<0.05) within the genotype. The upper letter in the bar not sharing the same letter differs significantly from each other. The genotype name illustrates Bt-1 (Bt Begun-1), Bt-2 (BARI Bt Begun-2), Bt-3 (BARI Bt Begun-3), and Bt-4 (BARI Bt Begun-4).

#### 3.3.2 Antioxidant enzymes

In our experiment, it has been seen that CAT activity was significantly increased and showed sensitivity in 100 mM and 150 mM concentrations of NaCl treatment compared to control (0 mM) in all genotypes. However, BARI Bt Begun-1 and BARI Bt Begun-2 showed better tolerance to salt treatment than other genotypes at 50 mM, 100 mM, and 150 mM concentrations. This is due to the fact that we discovered a larger quantity of CAT activity in these genotypes, which may help to ameliorate salt stress (Figure 8. A). We also measured the POD activity, and in our experiment, it was seen that POD activity significantly increased and showed sensitivity in 50 mM, 100 mM, and 150 mM concentrations of NaCl treatment compared to control (0 mM) in all genotypes (Figure 8. B). However, BARI Bt Begun-1 and BARI Bt Begun-2 demonstrated better tolerance to salt treatment at 50 mM, 100 mM, and 150 mM NaCl concentrations than the other genotypes.

**Figure 8.**
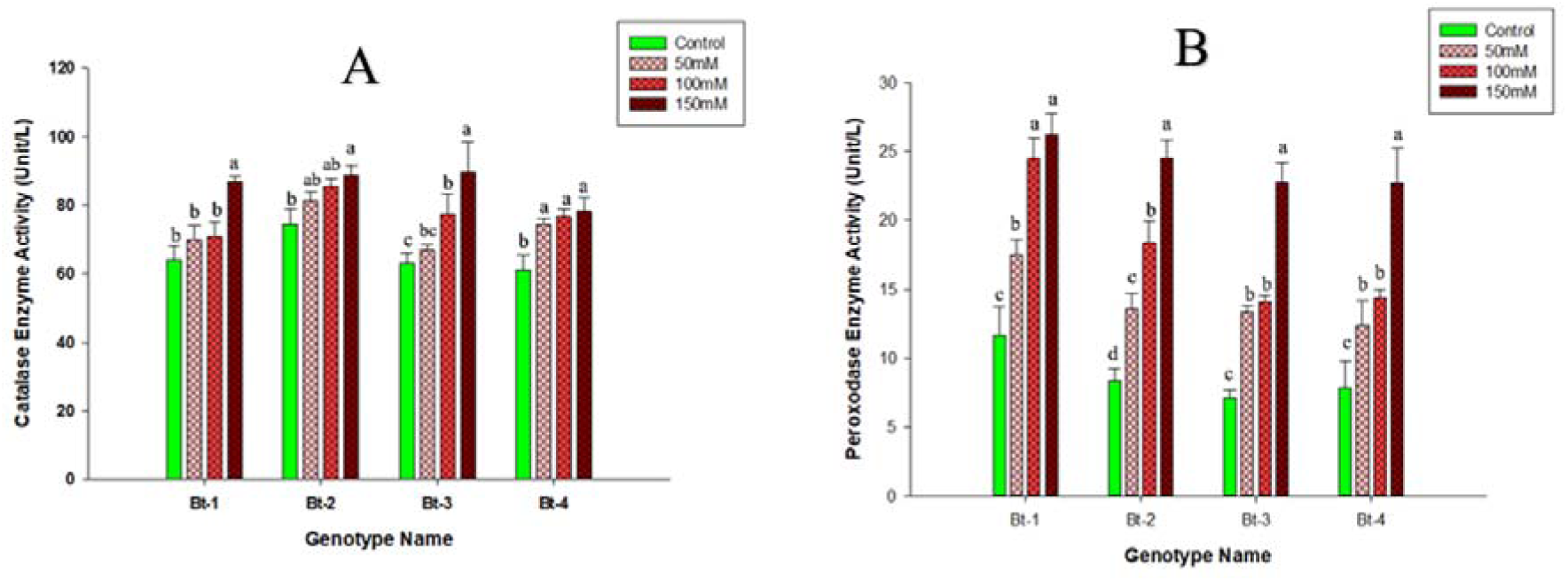
Catalase enzyme (CAT) activity and peroxidase enzyme (POD) activity of four BT Brinjal varieties in control and three different salinity levels (50 mM, 100 mM, and 150 mM salt concentrations). Y-axis represents the Catalase Enzyme Activity (Unit/L) (Figure 8. A) and Peroxidase Enzyme Activity (Unit/L) (Figure 8. B). Each bar indicates mean value ± standard errors (n=3). The upper letter in each bar indicates a pair-wise comparison (P<0.05) within the genotype. The upper letters in the bar, do not share the same letter is differ significantly from each other. The genotype name illustrates Bt-1 (Bt Begun-1), Bt-2 (BARI Bt Begun-2), Bt-3 (BARI Bt Begun-3), and Bt-4 (BARI Bt Begun-4).

#### 3.3.3 Proline content, anthocyanin content, lipid peroxidation, and hydrogen peroxide level

Proline content in all genotypes was significantly increased in 50 mM, 100 mM, and 150 mM NaCl concentrations compared to the control (0 mM) (Figure 9. A). As shown in Figure 9. A, BARI Bt Begun-1, and BARI Bt Begun-2 contain slightly more proline content than BARI Bt Begun-3 and BARI Bt Begun-4 to different stress treatments. As proline can mitigate the harmful effect of stress, BARI Bt Begun-1 and BARI Bt Begun-2 can be said to be more tolerant genotypes. Total leaf anthocyanin contents were also significantly increased and showed sensitivity in 100 mM and 150 mM concentrations of salt treatment compared to control (0 mM) in all genotypes (Figure 9. B).

**Figure 9.**
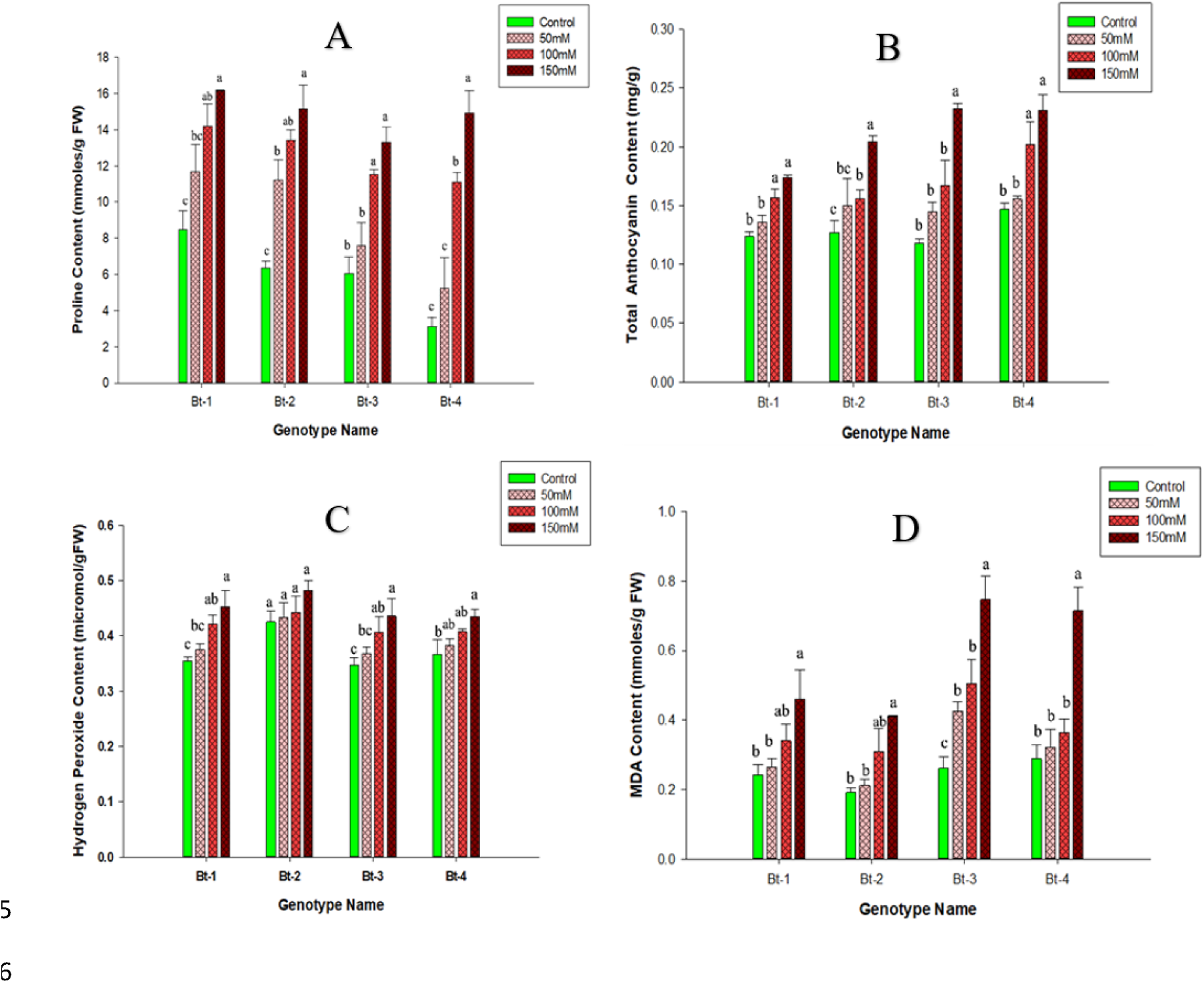
Proline content, Total anthocyanin content, Hydrogen peroxide content, and Malondialdehyde (MDA) content of four BT Brinjal varieties in control and three different salinity levels (50 mM, 100 mM, and 150 mM salt concentrations). Y-axis represents proline content (mmoles/g FW) (Figure 9. A), Anthocyanin Content (mg/g) (Figure 9. B), hydrogen peroxide content (micromol/gFW) (Figure 9. C), and MDA Content (m moles/g FW) (Figure 9.D), respectively. Each bar indicates mean value ± standard errors (n=3). The upper letter in each bar indicates a pair-wise comparison (P<0.05) within the genotype. The upper letters in the bar do not share the same letter is differ significantly from each other. The genotype name illustrates Bt-1 (Bt Begun-1), Bt-2 (BARI Bt Begun-2), Bt-3 (BARI Bt Begun-3), and Bt-4 (BARI Bt Begun-4).

Detection of H_2_O_2_ accumulation is critical, especially under abiotic stress conditions, because H_2_O_2_ is involved in oxidative cell damage and signaling mechanisms. We observed H_2_O_2_ content in all genotypes was also significantly increased only in 150 mM salt concentrations compared to the control (0 mM) (Figure 9. C). On the other hand, the lipid peroxidation level is represented as malondialdehyde (MDA) content. High MDA content indicates high oxidative damage in plants, while low MDA content indicates low damage induced by salt stress, which means more tolerance to stress. Usually, the MDA content increases noticeably upon exposure to salt stress. In our experiment, malondialdehyde (MDA) content was increased in all genotypes of 150 mM concentrations of salt treatment (Figure 9.D).But BARI Bt Begun-1 and BARI Bt Begun-2 showed lower increment trends of MDA content from control to 50 mM, 100 mM, and 150 mM concentrations of salt treatment (Figure 9.D).

## 4. Discussion

The current study was conducted to screen the tolerance levels of four Bt Brinjal varieties, namely BARI Bt Begun-1, BARI Bt Begun-2, BARI Bt Begun-3, and BARI Bt Begun-4, under different salt stress. Significant treatment, genotype, and their interaction under different levels of salt stresses showed a differential changes in phenotypic, biochemical, and antioxidant systems (Table 1, 2). A salt evaluation score (SES) was employed, and we have seen that BARI Bt Begun-1 and BARI Bt Begun-2 suffered less based on the SES score compared to BARI Bt Begun-3 and BARI Bt Begun-4 in 50 mM, 100 mM, and 150 mM concentrations of salt treatment (Figure 6. A) Similar findings were also observed for the number of leaves per plant. We have also seen that the decrease in shoot length was minimum in BARI Bt Begun-1 than in others in each stress treatment (Figure 4.B). Furthermore, the minimum decrement trends in root length were observed in BARI Bt Begun-1, and BARI Bt Begun-2 compared to BARI Bt Begun-3 and BARI Bt Begun-4 (Figure 4.A). Fresh weight is another important morphological parameter. The shoot fresh weight and root fresh weight were measured in our experiment. High salinity level causes reduced plant growth which ultimately leads to reduced shoot fresh weight. A similar response was observed in the current investigation. In our experiment, BARI Bt Begun-1 and BARI Bt Begun-2 varieties performed better in shoot and root fresh weight (Figure 5). We have also conducted dry weight measurements. In the case of the shoot dry weight, BARI Bt Begun-1 showed more tolerance to salt stress than the other three Bt Brinjal genotypes. Because the decrease in shoot dry weight was the smallest at various salt treatments in BARI Bt Begun-1 (Figure 5). Similar findings in other Brinjal genotypes were reported by [26] and [28].

We have also conducted many biochemical assays for characterizing four Bt Brinjal genotypes against salt treatment. It is stated earlier that increasing NaCl concentrations had increased the levels of proline content in the leaves of ’Adriatica’ and ’Black Beauty’, two Brinjal cultivars [17]. In our experiment, proline content was found to be increased after exposure to different salt stress treatments (Figure 9.A). Among them, BARI Bt Begun-1 and BARI Bt Begun-2 contained slightly more proline content than BARI Bt Begun-3 and BARI Bt Begun-4 to different stress treatments. As proline can mitigate the harmful effect of stress, BARI Bt Begun-1 and BARI Bt Begun-2 can be said to be more tolerant genotypes. The volatile aldehyde, like MDA, is a suitable marker for membrane lipid peroxidation and oxidative stress. The MDA levels were higher in water-stressed olive trees and in Coffea canephora [8]. In accordance with that the MDA content increased in the leaves of ’Adriatica’ and ’Black Beauty’, two Brinjal cultivars, upon exposure to increased NaCl concentrations [17]. The increment of lipid peroxidation (MDA content) was more in the BARI Bt Begun-3 and BARI Bt Begun-4 compared to Begun-1 and BARI Bt Begun-2 varieties (Figure 9.D), indicating that the Bt 1 and Bt 2 plants suffered less against stress treatments. On the other hand, the chlorophyll content was found to be decreased with an increasing salt concentration in all Bt Brinjal genotypes (7.A). In our experiment, total leaf chlorophyll contents decreased noticeably in all genotypes and showed sensitivity in 100 mM and 150 mM concentrations of salt treatment compared to the control. In contrast, the BARI Bt Begun-1 and BARI Bt Begun-2 genotypes were less sensitive to salt treatments at different salt concentrations. Carotenoid content was also decreased in all genotypes in all salt stress treatments, but BARI Bt Begun-1 and BARI Bt Begun-2 showed slight tolerance against salt stress. Similar findings were also described by [28] and [30] in other Brinjal genotypes. Environmental stressors cause a rise in antioxidant enzymes and metabolites, with their activity being relatively greater in stress-tolerant cultivars, indicating that increased antioxidant activity confers tolerance [19]. CAT and POD are two stress-responsive enzymes that are usually increased in plants to mitigate stress [7; 34]. In our study, it was seen that CAT activity was slightly increased and showed sensitivity in 100 mM and 150 mM concentrations of NaCl treatment compared to control in all genotypes (Figure 8. A). Among them, BARI Bt Begun-1 and BARI Bt Begun-2 showed greater tolerance than other genotypes in different concentrations of salt as a higher amount of CAT activity was observed in these genotypes, which can help to mitigate salt stress. Peroxidase enzyme (POD) activity was also significantly increased in 50 mM, 100 mM, and 150 mM concentrations of NaCl treatment compared to control in all genotypes (Figure 8. B). A higher amount of POD activity in BARI Bt Begun-1 and BARI Bt Begun-2 indicated a greater tolerance compared to other genotypes.

## 5. Conclusion

We successfully develop and established a cost-effective indigenous plastic containers-based hydroponics system and evaluated the salinity tolerance of the experimental Bt Brinjal genotypes in that system. The experimental results clearly indicated that Bt Brinjal varieties were severely affected by high salt concentrations i.e. 100 mM and 150 mM compared to the control. On the other hand, plants were less affected by 50 mM salt treatment compared to the control. Based on the measured traits, considerable genotypic differences were observed among the four Bt Brinjal varieties. However, BARI Bt Begun-1 and BARI Bt Begun-2, Brinjal varieties performed better compared to BARI Bt Begun-3 and BARI Bt Begun-4 against salt stress on the basis of genotypic performances. Thus, BARI Bt Begun-1 and BARI Bt Begun-2 varieties were ranked as moderately tolerant, and BARI Bt Begun-3 and BARI Bt Begun-4 varieties were classified as susceptible to salinity stress. Therefore, our results indicate that BARI Bt Begun-1 and BARI Bt Begun-2 varieties can be suitable for cultivation in saline-prone areas, which will improve the livelihoods of the poor marginal coastal farmers as well as secure the food supply. Further, the newly established hydroponic system could be used as an efficient, readily accessible, and cost-effective facility in modern research platforms.

