## Supplementary Figures for "Morpho-physiobiochemical dissection reveals insight into salt-induced differential responses in genetically modified Solanum melongena L. (Bt Brinjal) varieties using an indigenous hydroponic system"

Supplementary Figure 1. Seed germination and seedling establishment in hydroponic system

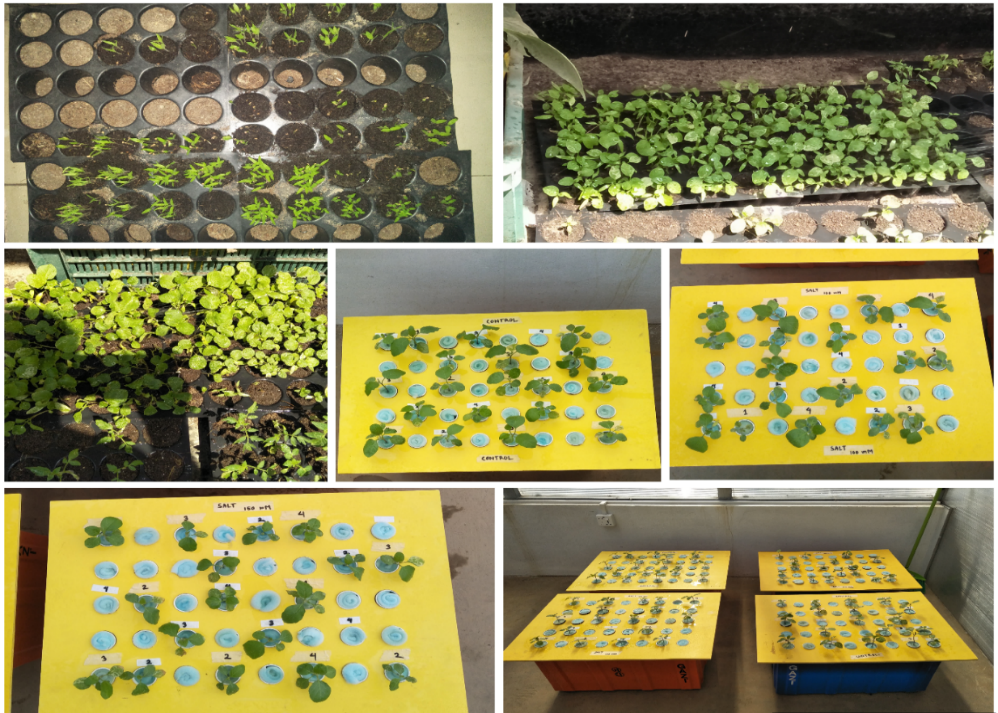

Supplementary Figure 2. Morphological differences between the treatments based on the

genotype

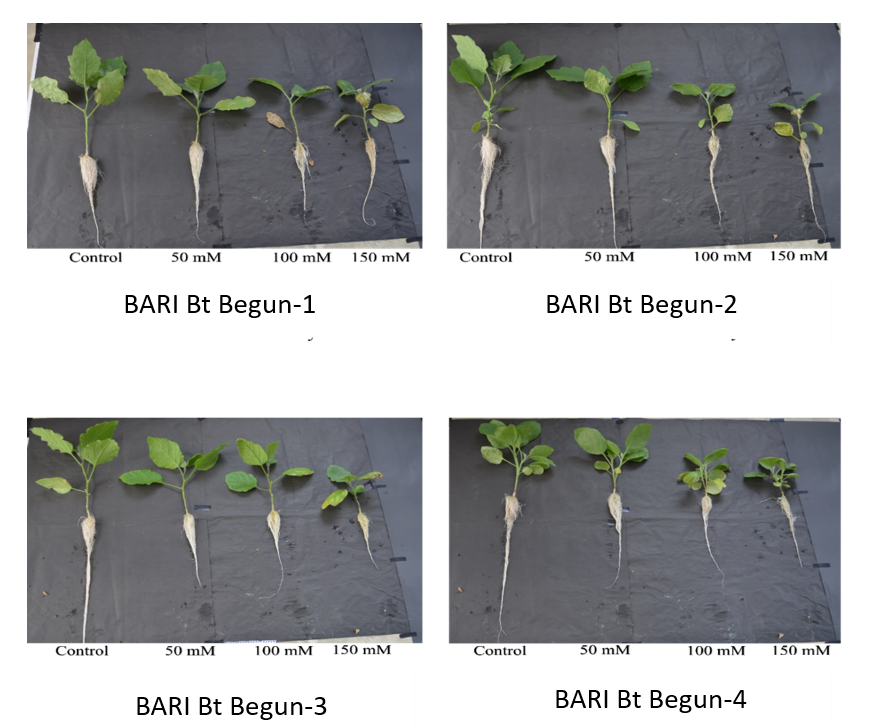

Supplementary Figure 3. Morphological differences among the genotypes based on the

treatments

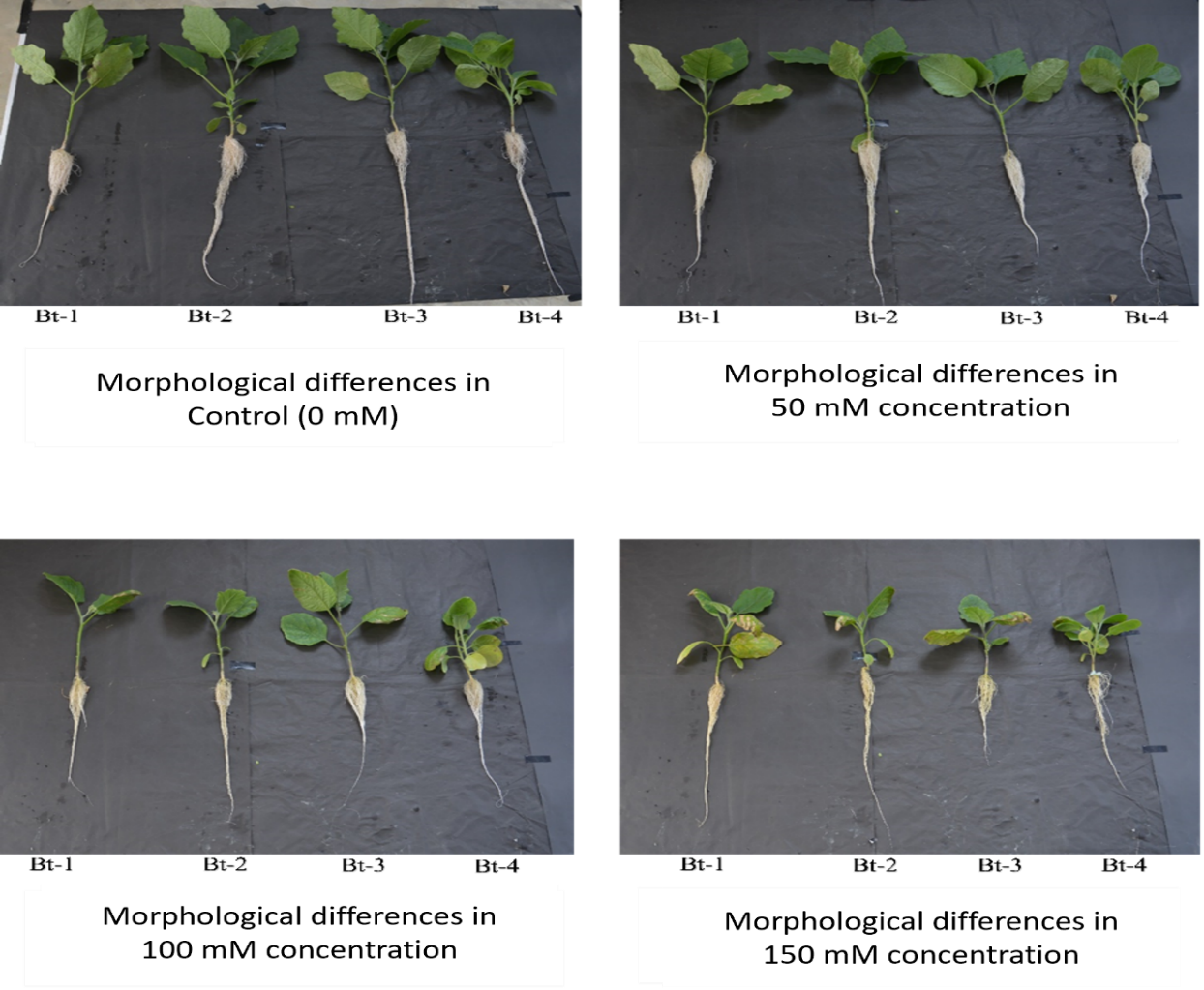
