## Supplementary Table for "Morpho-physiobiochemical dissection reveals insight into salt-induced differential responses in genetically modified Solanum melongena L. (Bt Brinjal) varieties using an indigenous hydroponic system"

Supplementary Table 1. Constitution of stock solution

| Stock | Chemical | Elements | (g)/1 liter |
| --- | --- | --- | --- |
| 1. | Ammonium Nitrate (NH_4_NO_3_) | N | 91.4 g |
| 2. | Sodium Phosphate monobasic Monohydrate (NaH_2_PO_4_.H_2_O) | P | 40.3 g |
| 3. | Potassium Sulfate (K_2_SO_4_) | K | 71.4 g |
| 4. | Calcium Chloride (CaCl_2_) | Ca | 88.6 g |
| 5. | Magnesium Sulfate heptahydrate (MgSO_4_.7H_2_O) | Mg | 324 g |
| 6. | Manganese Chloride Tetrahydrate (MnCl_2_.H_2_O) | Mn | 1.5 g |
|  | Ammonium Molybdate 4 Hydrate (NH_4_)_6_Mo_7_O_24_.4H_2_O] | Mo | 0.074 g |
|  | Boric Acid (H_3_BO_3_) | B | 0.934 g |
|  | Zinc Sulfate 7 Hydrate (ZnSO_4_.7H_2_O) | Zn | 0.035 g |
|  | Copper Sulfate 5 Hydrate (CuSO_4_.5H_2_O) | Cu | 0.031 g |
|  | Fe-Na-EDTA | Fe | 10.4 g |
